# Larger and polymorphic noctuid moths tend to show less inter-annual abundance variation in the canopy of a temperate forest

**DOI:** 10.1101/2025.03.13.643003

**Authors:** Freerk Molleman, Roman Wąsala, Ahmet Tambay, Robert B. Davis, Erki Õunap, Urszula Walczak

## Abstract

Inter-annual variation in insect abundance and seasonal phenology can be related to species traits such as body size, larval diet, overwintering stage, and colour variations. We sampled noctuid moths in the canopy of a forest in Western Poland using flight-interception traps during two vegetative seasons. We calculated inter-annual variation for 31 species, and date of peak abundance and length of the flight season for 18 species. We found that for the 23 species for which we had phylogenetic information, larger moths and those with adult colour variation showed less inter-annual abundance variation, which corroborates results of previous studies. We found no indication that phenological traits are associated with the tested species traits. However, species that can feed on the dominant broad-leaved tree in which the traps were placed (*Quercus petraea*) tended to have a later date of peak abundance than other species. To draw more robust conclusions, future research should encompass a longer time span and a broader range of species.

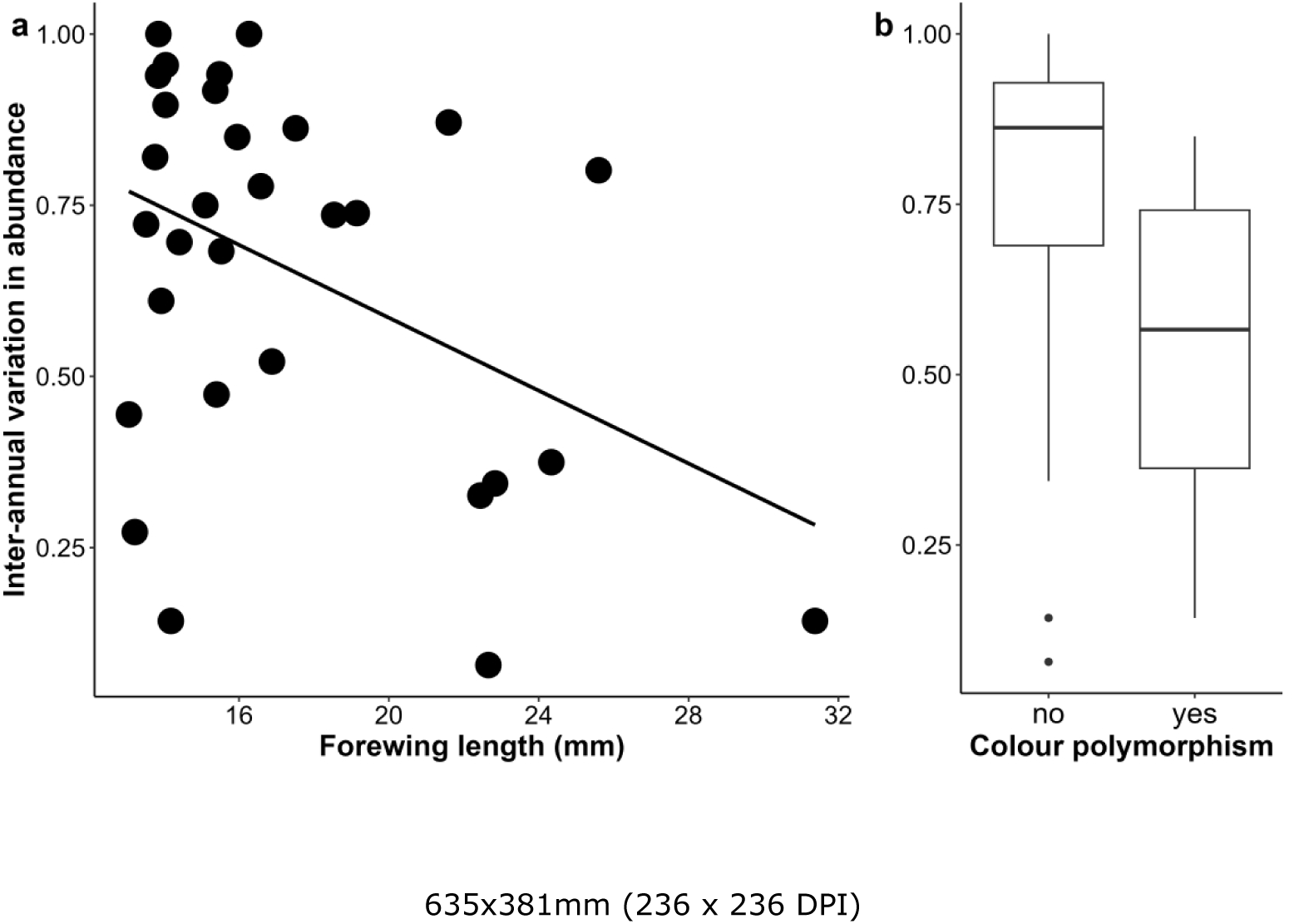

## INTRODUCTION

Climate change and other anthropogenic factors are impacting both the population sizes and phenology of insects in temperate habitats (Wagner et al., 2021, Fox et al., 2014), but we have limited understanding on how this is affected by species traits. The abundance of adult insects at a given location varies as a result of their phenology, population dynamics, and movements (Molleman, 2018). These processes are, on the one hand, affected by environmental variation (in particular weather), and on the other, by species traits such as diet and body size (Seifert et al., 2023, Tammaru and Haukioja, 1996, Gaston, 1988). Moreover, traits of a species can influence how it is affected by environmental variation (Végvári et al., 2015). For example, noctuid moths overwintering as adults show greater phenological shifts in response to climate change than those overwintering as other life stages (Végvári et al., 2015). Therefore, more insight into insect temporal abundance variation and its drivers is needed to predict species’ responses to the changing climate (Coulthard et al., 2019).

Moths, in particular noctuids, are often used as a model group to study these temporal dynamics, as they are often abundant, species-rich, and can be sampled with light traps at ground level. As the effectiveness of light traps is affected by ambient light (moon phase, cloud cover) and vegetation structure, and such traps attract moths from variable distances, passive traps such as Malaise traps can potentially provide more reliable data (Butler et al., 1999, Busse et al., 2022, La Cava et al., 2024, Merckx and Slade, 2014, Truxa and Fiedler, 2012). Furthermore, studies including trapping in the canopy have recovered pronounced vertical stratification, so that parts of the moth community are missed by ground-level sampling (De Smedt et al., 2019, Ashton et al., 2016, Ruchin, 2023, La Cava et al., 2024, Böttger et al., 2025, Fröhlich et al., 2007). However, analyses of temporal patterns in the abundance of moths found in the canopy of forests are scarce, even for the temperate zone.

Variation in inter-annual abundance of adult moths results from population dynamics within the typical dispersal range of the species. Thus, the temporal patterns observed at a particular location will, in some species, represent very local population dynamics (possibly even at the level of an individual tree), and in highly dispersive or migratory species may reflect those happening over hundreds of kilometres. Thus, abundance fluctuations are not the same as population dynamics, and this may lead to wrongful interpretation of abundance fluctuations measured at a given location (Molleman, 2018). Species traits that may affect abundance fluctuations are often correlated with each other. Migratory behaviour tends to be more common in larger-bodied species and in those that are more polyphagous (Nieminen et al., 1999, Logghe et al., 2024). In addition, higher wing loading (large wings relative to body mass) may allow longer and faster flights, which are suitable for long-distance dispersal or migration, whereas lower wing loadings are associated with slower flights and hovering (Betts and Wootton, 1988). While studies have shown that dispersal traits affect population trends and range shifts, their relationships with the degree of inter-annual variation remain largely unexplored (Jones, 2014). The effect of adverse weather on adults may be stronger for smaller-bodied and shorter-lived species, and individuals of smaller-bodied moth species also tend to be shorter-lived (Holm et al., 2016). In particular, smaller-bodied species may more often fail to reproduce when their short life span happens to coincide with adverse weather, possibly leading to more erratic population fluctuations (Gaston, 1988, Gaston and Lawton, 1988). Consistently, larger arthropods feeding on bracken ferns were shown to have less variable abundance than smaller-bodied species (Gaston and Lawton, 1984). Overall, body size is likely correlated with the degree of inter-annual variation in abundance due to such indirect relationships (Gaston and McArdle, 1994, Gaston and Reavey, 2009, Jones, 2014).

Larval diet breadth can also affect abundance fluctuations. When population dynamics result from interactions with a small number of resources (host-plant species) and natural enemies (parasitoids, predators, and pathogens), the overall population trends will tend to be more stochastic (MacArthur, 1955). In contrast, long-term studies on moths usually show more inter-annual abundance variation in species with more polyphagous larvae, which may have to do with the habitat, as polyphagous species are often associated with early successional vegetation (Spitzer and Jaroš, 2009 and associated bibliography). Species that require close synchrony between their phenology, the phenology of host plants, and suitable weather, also tend to show high levels of inter-annual abundance variation (Price, 2003, Tammaru and Haukioja, 1996). Specifically, populations of spring feeders often appear to respond strongly to variation in synchrony between egg hatching and budburst phenology (e.g. winter moths) and also suffer more from unseasonably late episodes of frost than those that breed during the summer (Hunter and Elkinton, 2000, Varley and Gradwell, 1967). Lastly, variable adult colouration (polymorphism), which is thought to reduce predation risk when moths occur at high densities, is associated with reduced inter-annual variation in the abundance of noctuid moths (Forsman et al., 2015). Overall, we can expect that body size, wing loading, host-plant relationships, and adult colour variation are correlated with inter-annual variation in moth abundance.

Moth phenology is usually controlled by temperature, photoperiod, or a combination of these factors (Valtonen et al., 2011), and is thus likely to both plastically and adaptively change as the climate is warming (Teder, 2020, O’Neill et al., 2012, Hickinbotham et al., 2024, Hällfors et al., 2021). Other species traits are also correlated with phenology and may affect how phenology responds to climate change. This may include body size (Foerster et al., 2024). For example, smaller-bodied, spring-feeding species may be expected to complete their development faster and thus emerge earlier in the season. Across the entire season however, species of geometrid moth whose larvae develop in the spring are generally larger than species developing during late summer (Seifert et al., 2023). Overwintering stage is also related to moth phenology, as species overwintering as pupae emerge earlier than those that overwinter as larvae or eggs (Hickinbotham et al., 2024). Furthermore, species feeding on woody plants tend to emerge later than those feeding on herbaceous plants (Hickinbotham et al., 2024). Moths feeding on woody plants also show larger shifts in flight periods but smaller increases in voltinism in response to climate change than those feeding on herbaceous plants (Altermatt, 2010). Moreover, phenological responses to climatic variation can be stronger for species with later overwintering life stages (e.g. overwintering as adults vs as eggs; Hickinbotham et al., 2024). Similarly, spring-flying species shift their phenology more in response to climate warming than summer- or autumn-flying species (Maurer et al., 2018). Overall, body size, timing of development, overwintering stage, and type of host plant can be expected to correlate with the phenology of moths.

To preliminarily gauge how species traits may be related to inter-annual variation in adult abundance and phenology of moths in the canopy of a temperate forest, we analysed data on noctuid moths in the canopy of a forest in Western Poland that were caught using flight-interception traps during two years. To our knowledge, this is the first such study performed on noctuids in the canopy of a forest, where a distinct fauna may be encountered. Moreover, we used traps that do not attract but only intercept ambient moths, in contrast to most moth studies that use light traps (see e.g. Butler et al., 1999, Busse et al., 2022 for moth sampling with Malaise traps).

## METHODS

### Study site and insect collection

We sampled moths in Puszcza Zielonka forest, Western Poland (52.524, 17.059). Most compartments of this forest are dominated by pine (*Pinus sylvestris* L.) or sessile oak (*Quercus petraea* (Matt.) Liebl.), while some smaller areas have mainly beech (*Fagus sylvatica* L.), pedunculate oak (*Q. robur* L.), or spruce (*Picea abies* (L.) H. Karst.; Forest_Data_Bank, 2018). In some areas, there is a subcanopy of hornbeams (*Carpinus betulus* L.; Forest_Data_Bank, 2018).

We trapped insects with flight-interception traps (Figure 1; Gossner et al., 2014) during the vegetative seasons of 2019 (April–October) and 2020 (March–October) in the canopy of 34 sessile oak trees (the same trees as used in Molleman et al., 2022 with some additions). This study was designed to study the insect fauna associated with individual trees in a forest, and effects of tree neighbourhood and tree traits on noctuoid moth communities captured will be treated elsewhere. Focal oak trees were located in four stands near Zielonka village and seven stands near Kamińsko village in sets of two to four trees (Figure 2). Within sets, trees were selected to vary widely in the species identity of neighbouring trees (mainly oaks, pines, beeches, or hornbeams). The traps had both a bottom collector, and a top collector functioning like a Malaise trap. They were suspended over high branches within the canopy at a height of 15–25 metres. Traps were serviced every 3–4 weeks. Usually, not all traps could be serviced on one day, causing slight differences in collection dates among trees. Insects were preserved in the traps in polypropylene glycol and then transferred to jars with 70% ethanol, and these were kept at -20 °C. The macro moths were sorted from the samples and provisionally set (curated at AMU). Moths were identified to species using genitalia dissection when needed (Fibiger, 1990, Fibiger, 1993, Fibiger, 1997, Fibiger and Hacker, 2007, Fibiger et al., 2009, Fibiger et al., 2010, Ronkay and Ronkay, 1994, Ronkay and Ronkay, 1995, Ronkay et al., 2011, Ronkay et al., 2001).

**Figure 1.**
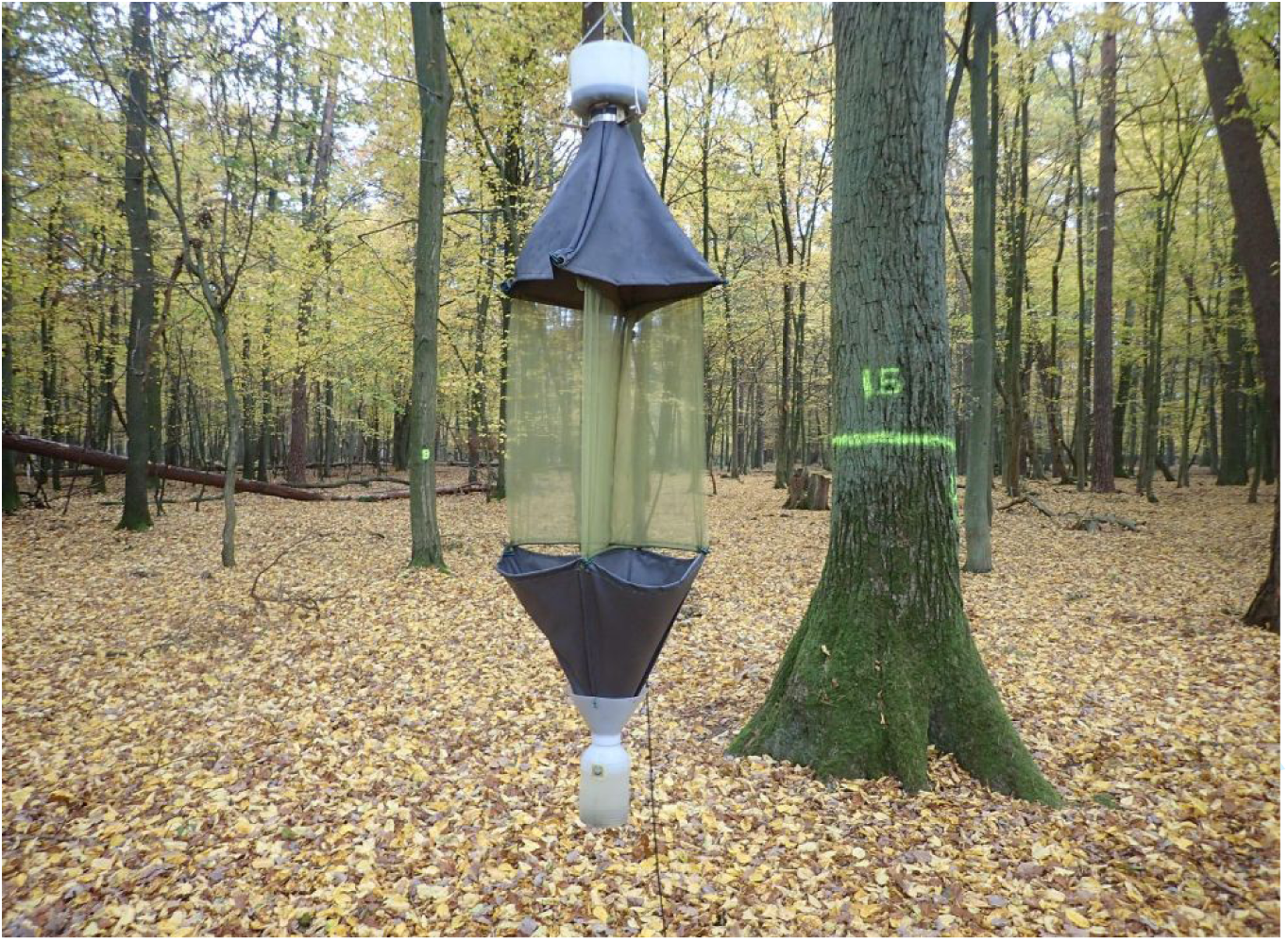
Flight-interception trap suspended on a rope that passes over a high branch. Traps were pulled into the canopy using a pulley system and lowered for servicing about once a month. Insects intercepted by the see-through mesh either climbed up and ended up in the top container or fell into the bottom container. Both containers contained polypropylene glycol to preserve the insects. The bottom container was fitted with a hole covered with fine wire mesh near the top, to allow excess rainwater to flow out while retaining captured insects.

**Figure 2.**
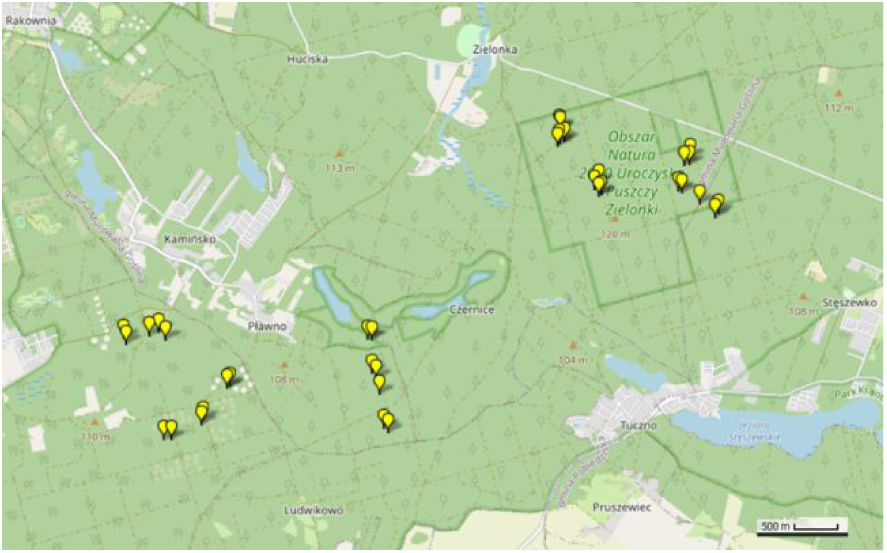
Location of flight-interception traps in Puszcza Zielonka, Poland. Map created by GPSVisualizer.

### Species traits

To obtain body size data, we measured the forewing length and thorax width of up to ten individuals of each species. We used specimens from our traps whenever possible, but completed the data using other specimens collected in the region. We then calculated wing loading as wing length^2^/thorax width^3^. To obtain information on larval diet, overwintering stage, feeding period, and migratory behaviour, we used authoritative websites (Wagner, 2024, Lepiforum, 2020, Jonko, 2020). Polymorphism was defined as the presence of multiple, discrete variants (or morphs) within a single population, the rarest of which is too common to be solely due to recurrent mutation (McLean and Stuart-Fox, 2014, Gray and McKinnon, 2007, Ford, 1945). Our assessment of polymorphism was based on our own catches of moths, mainly in Central Europe, and reflects the range of variability documented in the literature for the entire European range of the species (Lepiforum, 2020). A prominent example in our sample is the large yellow underwing *Noctua pronuba* (Linnaeus, 1758), which exhibits a wide range of colour forms (Cook and Sarsam, 1981). Another example is the dark arches *Apamea monoglypha* (Hufnagel, 1766), which has light and dark colour forms (Askew et al., 1971). Larval diet was parsed in four ways. First, we distinguished species by their ability to feed on oak trees (the tree species in which traps were hung) as specialist of oak, generalist including oak, or non-oak feeding. Secondly, we classified moths as specialists, oligophagous, generalists, or grass-feeding. Third, we classified them by growth form (tree, herb, tree & herb, grass). Finally, we classified them based on food type: leaves, roots, or stems). Overwintering stage was classified as egg, larva, pupa, or adult. Larval feeding period was classified as spring, summer, autumn, and combinations thereof, or the entire growth season. Whether a species is known to migrate or to have extensive adult colour variation within populations was coded as yes or no.

### Data analysis

To calculate inter-annual abundance variation, we first summed the number of individuals caught for each species for both years and for the two study years separately. For each species for which we caught at least ten specimens, we then calculated inter-annual variation as ‘one minus (the lowest abundance divided by the highest abundance)’. Given that we have two years of data, this is preferable over the coefficient of variation (CV) recommended for larger data sets (Gaston and McArdle, 1994), as it is simple, little affected by the mean, and does not rely on assumptions on the data distribution.

We reconstructed the phenology of each species of moth collected from the traps in several steps. The recorded collection date of an individual was when the trap was serviced (servicing day), and it could thus have been trapped in the 3–5 week period (trapping period) since the previous servicing of the trap. It could take multiple days to collect material from all the traps, and the servicing day was defined as the middle of this period. To calculate the number of individuals of a given species trapped per day, we first summed the number of individuals over all the traps in a given trapping period, and then divided this by the length of the trapping period. To calculate the abundance per trap per day, we then divided this by the number of traps that were active. Assuming homogenous capture probability during each trapping period, the middle of the trapping period was taken as the date associated with the abundance per trap per day. For the species of which we caught at least 50 individuals, we then plotted the abundance per trap per day against date for both years separately and fitted a line using the loess function in R (R_Core_Team, 2025). From this line, we extracted the date of peak abundance and the first and last occurrence of the moth species (abundance > 0). The length of the flight period was then calculated as the time between the first and last occurrence of the species.

We created a timed phylogenetic tree of Noctuidae to be able to perform phylogenetic comparative analyses, relying on the recent phylogeny of Noctuidae (Nedumpally et al., in prep). For some species, we lacked the data to include them in the phylogeny. *Agrotis segetum* was represented in the phylogeny by another species from the same genus for which phylogenetic information was available, *A. puta*. We used this phylogeny as the fixed topology and incorporated the corresponding molecular data matrix. Since the full molecular dataset was too large for node age calculations, we selected the 24 longest protein-coding loci that contained data for at least 90% of the taxa, totalling 53,990 bp. First, we converted the initial tree into an ultrametric one using the makeChronosCalib and chronos functions in the R package ape (Paradis and Schliep, 2019). Because the full tree of 333 species was too large for node age calculations, we split the tree and corresponding molecular dataset into two subsets. We used the drop.tip function in ape to exclude almost all Noctuinae: Xylenini + Apameini from one subset (subset 1) and almost all Noctuinae: Noctuini + Hadenini s.l. from the other (subset 2). Both subsets retained 37 outgroup species from eight families of Macroheterocera (Drepanidae, Uraniidae, Geometridae, Lasiocampidae, Saturniidae, Sphingidae, Notodontidae, and Erebidae) along with 90 to 91 species of Noctuidae. The final data matrix included 246 species for subset 1 and 227 species for subset 2. To determine the optimal partitioning scheme based on predefined loci, we used MODELFINDER (Kalyaanamoorthy et al., 2017) within IQ-TREE 2.3.2 (Nguyen et al., 2015) on the CIPRES portal (Miller et al., 2010). We followed the results suggested by MODELFINDER when dividing the data into partitions for node age calculations, resulting in 13 partitions for subset 1 and 15 for subset 2.

We calculated node ages using BEAST v1.10.4 (Suchard et al., 2018) at the High-Performance Computing Center of the University of Tartu (Anonymous, 2025). Since test runs with more complex clock models failed to converge, we applied a strict molecular clock, linked clock and tree priors across all partitions, and implemented a birth–death tree prior with incomplete sampling (Stadler, 2009) for the final runs. To time-calibrate both ultrametric trees, we defined five calibration points for each subset based on Kawahara et al. (2019), with ages (in millions of years) as follows: root of the tree (Normal distribution, mean 93.7 ± 11.1 SD), Noctuoidea (Normal distribution, mean 77.6 ± 11.0 SD), Erebidae + Noctuidae clade (Normal distribution, 68.6 ± 10.2 SD), *Chloridea* + *Helicoverpa* clade (Normal distribution, 9.3 ± 5.6 SD), and *Spodoptera* (Normal distribution, mean 10.6 ± 5.7 SD). For each subset, we ran a single BEAST analysis with Bayesian MCMC for 30 million generations, sampling every 1000th generation. After inspecting the results in TRACER v1.7.2 (supplementary software to BEAST), we discarded the first 3 million generations as ‘burn-in.’ We used TREEANNOTATOR v1.10.4 (also supplementary software to BEAST) to construct the final trees for both subsets. Finally, we combined the two partitions to generate the full tree, making minor branch length adjustments in the basal branches to ensure ultrametricity.

To investigate relationships between species traits and inter-annual abundance variation, we conducted phylogenetic comparative analyses implementing a series of single-predictor models. For continuous predictors, such as mean forewing length, we performed PGLS analyses using the R package ape (Paradis and Schliep, 2019). When possible, we applied the Pagel model to account for phylogenetic signal. We tested residual normality using a Shapiro– Wilk test and QQ plot inspections. When necessary, we reran PGLS analyses after removing outlying data points; specifically, values that clearly deviated from the theoretical normal line in QQ plots were considered outliers. For categorical predictors, such as food type, we conducted a phylogenetic ANOVA using the phylANOVA function in the R package phytools (Revell, 2012). For categorical predictors that included groups with only one species, we ran analyses both with and without those groups, as phylANOVA only provides pairwise t-values for groups with more than one species.

We first tested if the phenology differed between the two study years using paired t-tests. To explore relationships between species traits and phenology, we first made a correlogram for each study year using the R package corrplot for traits that were continuous or binary (Wei and Simko, 2021). As the number of species with at least 50 individuals captured was too small to implement phylogenetic comparative methods, we then implemented linear models for the average date of peak abundance and the average length of the flight period with as predictors the species traits in R (R_Core_Team, 2025). As dependent variables, the full models had either average peak date or length of the flight season, and as predictors mean forewing length, thorax width, wing loading, diet (oak specialists, generalist including oak, non-oak feeding), food type, plant form, overwintering stage, feeding period, variation in adult coloration, and whether it is known to migrate. We then performed AICc model selection using the R package MuMln (Barton, 2022), limiting the maximum number of predictors to three to avoid over-parametrization. We implemented selected models and determined if model assumptions were met by plotting the models and particularly inspecting QQ plots.

## RESULTS

### Overview of the data

We caught 5300 individual noctuoid moths (link to Dryad data, see additional files). For 31 species, we caught at least ten individuals (Appendix 1), and for 23 of these we also had phylogenetic information. For 18 species, we captured at least 50 individuals (Figure 3).

**Figure 3.**
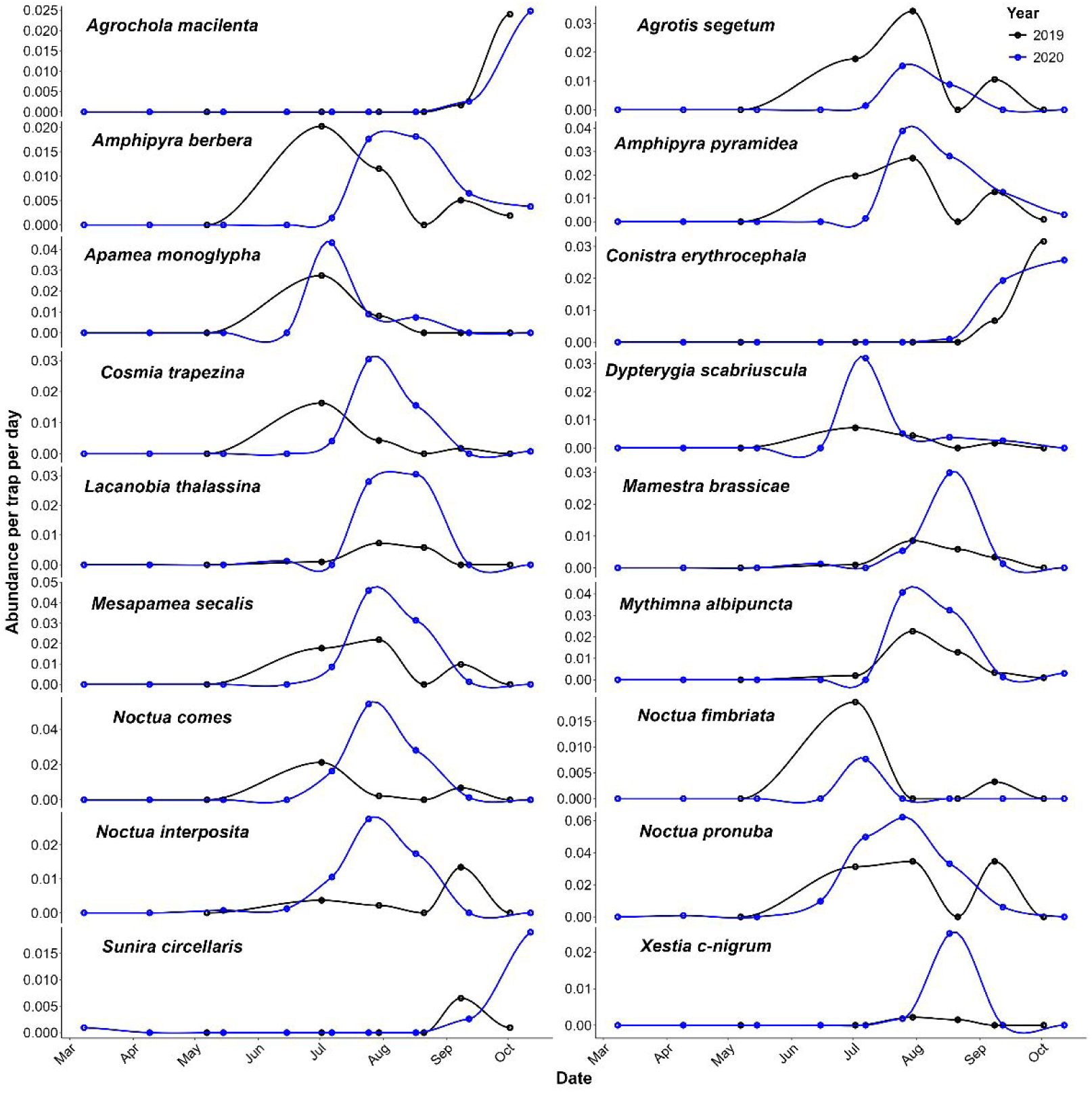
Phenology of the most abundant noctuid moths (N > 50) during 2019 (black) and 2020 (blue) sampled with flight-interception traps in the canopy of a forest in Western Poland.

### Phylogeny of the study species

The sufficiently abundant noctuoids sampled (N > 10) all belonged to the family Noctuidae. We reconstructed the phylogeny of the species for which we had phylogenetic information (Figure 4).

**Figure 4.**
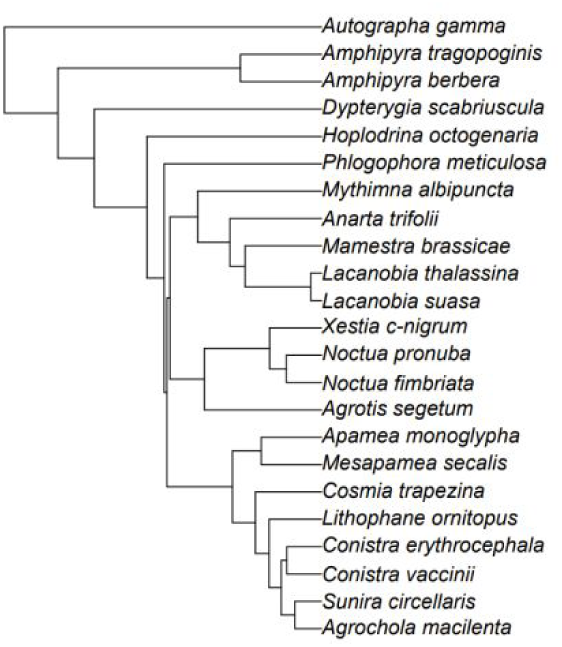
Time-calibrated phylogeny of the noctuid moths used in this study.

### Inter-annual variation in abundance

Inter-annual variation in abundance varied from species that were caught only during one of the study years, to species that were quite evenly distributed across the two years (Figure 5a). Inter-annual variation in abundance tended to be less pronounced in noctuid moths with longer wings (Figure 5a, Table 1), and in species with pronounced adult colour variation (Figure 5b; Table 1). However, after correction for multiple testing, these relationships were not significant (Table 1). Other species traits did not show significant relationships with inter-annual variation in abundance (Table 1). Removal of outliers had no qualitative effect on the results.

**Fig 5.**
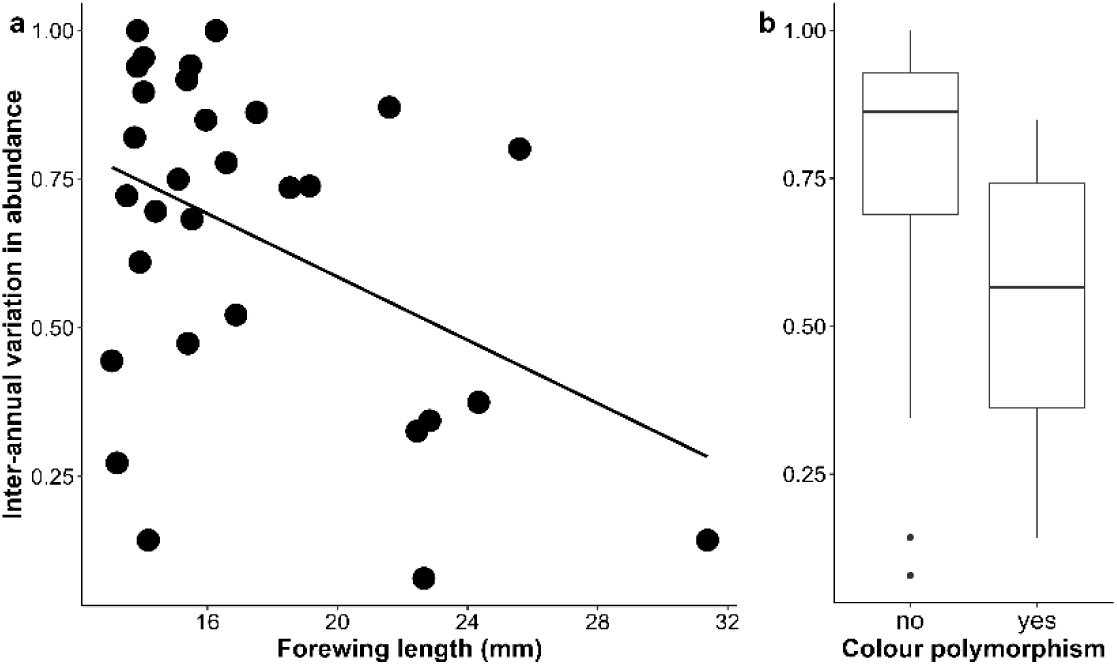
Relationship between inter-annual variation in adult abundance of noctuid moths and a) average forewing length, and b) adult colour polymorphism. Variation in abundance can vary between zero (the same abundance in both years, and one (only caught during one of the study years). See Table 1 for results of statistical analyses accounting for phylogeny.

**Table 1.**
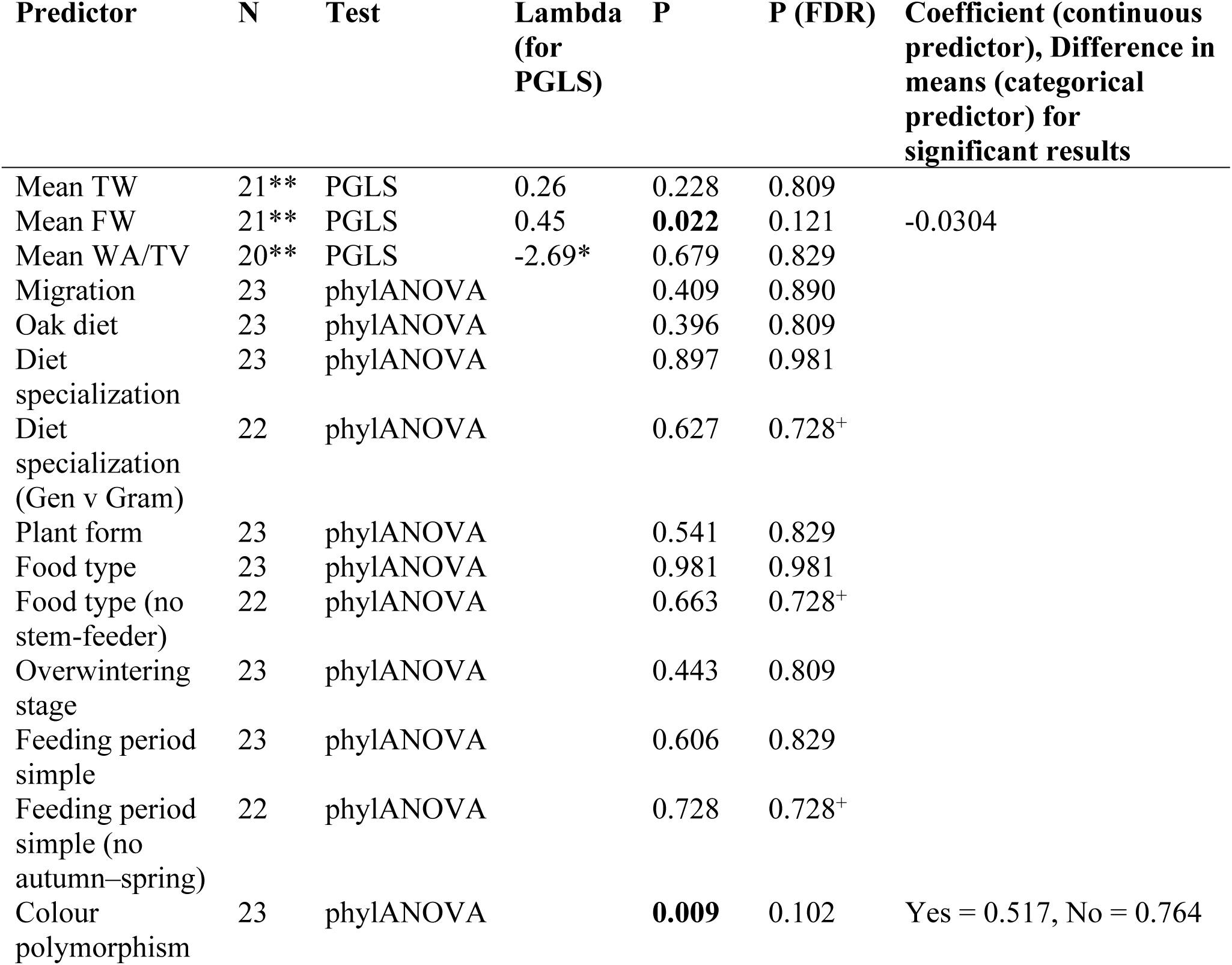
Results of phylogenetic comparative analyses of inter-annual variation in noctuid moth abundance. Where N is marked **, outliers have been removed. The exclusion of outliers had no qualitative impact on the result of the analysis. Where lambda is marked *, this indicates a suspicious lambda estimate; the analysis using mean WA/TV as a predictor was already problematic before residual outliers were removed and could not be run using the corPagel function in ape (R). Using the corBrownian function after outliers are removed also produces a non-significant result. P (FDR) values relate to a test including p-values from single-predictor tests where categorical predictors included all states. Where p (FDR) is marked +, this indicates a value for a test rerun removing states where n = 1. Rerunning the test in this manner had no qualitative impact on other p (FDR) values. The exclusion of outliers had no qualitative impact on the result of the analyses. P-values < 0.05 are in bold font. Thorax width = the average width of the thorax; Wing length = average forewing length; WA/TV = wing loading calculated as Wing length^2^/Thorax width^3^; Oak diet = whether the larval diet includes the dominant broad-leaved tree species (*Q. petraea*) in which traps were placed; Dietary specialization = specialist (one host plant genus), generalist (feeding on multiple genera, or grass-feeding); Plant form = whether larvae feed on trees, herbs, trees and herbs, or on grasses; Food type = leaves, roots, or stems; Food type excluding stem = same as Food type but excluding the one species feeding on stems; Overwintering stage = egg, larva, pupa, imago, or not overwintering in Poland; Feeding period = time of year when larvae are feeding (spring, summer, spring & summer, spring & autumn, summer & spring, vegetation period); Colour polymorphism = whether the species is known to have adult colour variations within the population; Migration = whether the species is known to migrate or have mass dispersal events.

### Phenology

The peak abundance of the moths ranged from June to October, and the flight period tended to be long, ranging from 123 to 224 days (Figure 3). Across these 18 species, the date of peak abundance did not differ significantly between the two study years (paired t-test: t = -1.66, df = 17, p = 0.12) nor did the length of the flight season (t = 0.34, df = 17, p = 0.74). Oak-feeding species tended to have a later peak date than non-oak-feeding species (significant at p = 0.01 in 2020, Figure 6). For 2020, overall abundance appeared associated with the length of the flight season (p = 0.06, Figure 6), which may indicate that at small sample sizes, the length of the flight season may be underestimated, but this was not the case in 2019 (Figure 6).

**Figure 6.**
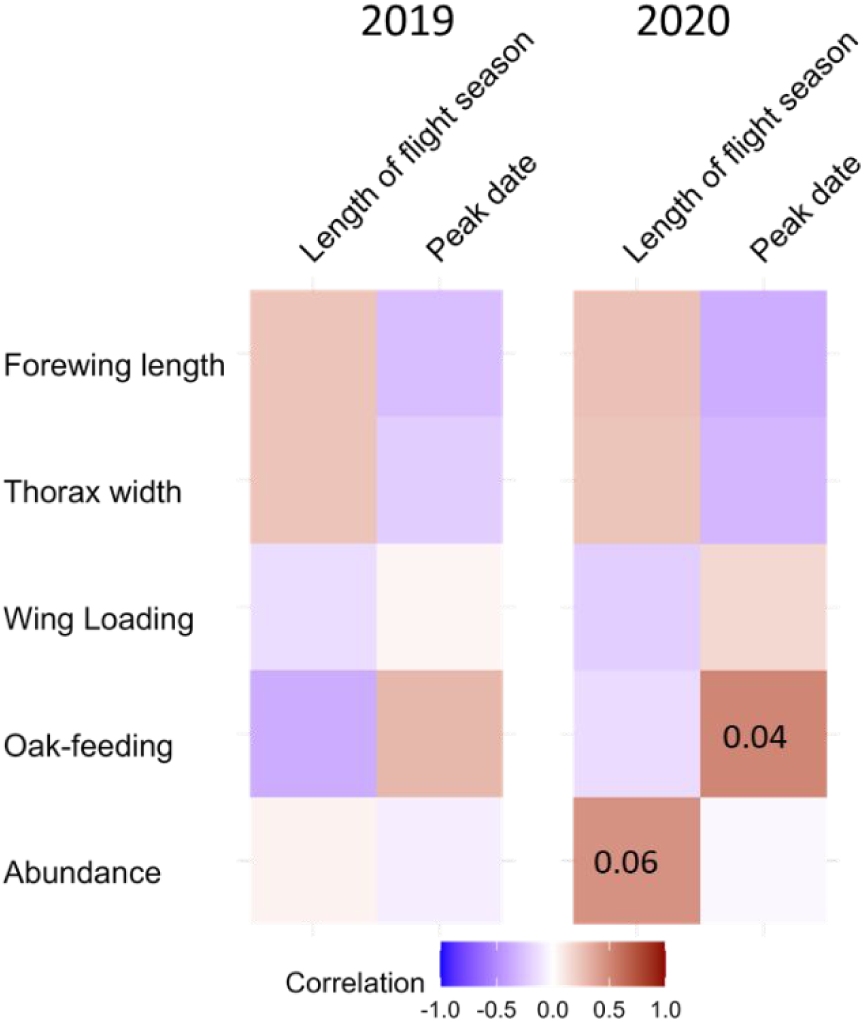
Correlations between phenology and species traits for 2019 and 2020 with p-values when they were < 0.1.

Model selection based on AICc for the average length of the flight season resulted in a model without predictors, followed by a model with forewing length. However, forewing length was not a significant predictor (R2 = 8%, Adj-R2 = 2%, F = 1.38 on 1 and 15 DF, p = 0.26). Similarly, the top model for the average date of peak abundance did not include any predictors, and the second best included only wing length, but this was not significant either (R2 = 11%, AdjR2 = 5%, F = 1.81 on 1 and 15 DF, p = 0.20). Nevertheless, similar to the correlations (Figure 6), the average peak date was significantly later for oak-feeding moths (R2 = 24%, AdjR2 = 20%, estimate = -29.58, F = 2.268, p = 0.036).

## DISCUSSION

We sampled noctuid moths in a forest in Poland during two seasons and explored relationships between some species traits that have been suggested to correlate with inter-annual variation in abundance or phenology. We used flight-interception traps rather than the usual light traps, avoiding some of the biases associated with the latter. Furthermore, we trapped in the canopy in contrast to other multi-year moth studies that were all performed at ground level. Even though the two-year duration is the minimum for looking into inter-annual variation, and the number of species for which we had enough data for comparative analyses was only 23, we found that larger moths and those with adult colour variation showed less abundance variation, which corroborates results of previous studies (e. g. Gaston and Lawton, 1988, Forsman et al., 2015). Notably, species traits with more than two levels never showed a significant effect, which may be due to the low number of species available in our dataset. In addition, we described the phenology of 18 species but found no indication that phenological traits are associated with the tested species traits. However, species that were able to feed on the dominant broad-leaved tree in which the traps were placed (*Q. petraea*) tended to have a later date of peak abundance than other species.

Admittedly, the duration was short (only two years) and the number of species was small for comparative analyses. Moreover, our results must be treated with caution as no relationship remained significant after correction for multiple testing. To address if abundance fluctuations differ between canopy and understory species, future studies should sample simultaneously in both strata for extended periods. Further studies on phenology should also take into account voltinism, as some species may have multiple peaks per year or overlapping generations.

The effect of body size on inter-annual variation in moth abundance can result from various indirect effects (see discussion in Gaston and McArdle, 1994, and Introduction). It may be related to life span, dispersal, and susceptibility to weather. Larger moths tend to live on average longer, so that they are less affected by short-term weather events that may prevent shorter-lived species from completing their life cycle. They may also be better dispersers, evening out local population dynamics over a wider area. Finally, larger individuals may be more robust to adverse weather. Colour polymorphism is thought to allow populations to maintain higher densities, as it reduces search image formation in predators such as birds (Bond, 2007, Karpestam et al., 2016). The association between reduced inter-annual variation in abundance and colour polymorphism that we found in our data, was shown before for noctuids using longer time series of more species (Forsman et al., 2015). Overall, our results suggest that small-bodied species lacking adult colour variation may vary more in abundance and may thus be more prone to (local) extinction. Further work may uncover whether this translates into greater susceptibility to climate change and other stressors.

That species that are able to feed on the dominant broad-leaved tree in which the traps were placed (*Q. petraea*) tended to have a later peak date than other species, suggests that these moths eclose later in the season than other species that developed in the understory or immigrated. Perhaps the late budburst of oaks makes their leaves available for herbivores later than those of most other plant species (Molleman and Walczak, 2024, Wesołowski and Rowiński, 2006), thus causing their later emergence. This would be similar to the emergence of European grape vine moths depending on grape variety (Thiéry et al., 2014). Otherwise, no species traits appeared associated with their date of peak abundance, nor with the length of their flight season. The results of these analyses, however, must be taken with some caution, as we had only 18 species with at least 50 specimens, and we were not able to take into account their phylogeny.

### Conclusions

This is the first study to document temporal abundance fluctuations of noctuid moths using: a) flight-interception traps, b) in the canopy of a temperate forest, and c) in Poland. Future studies could sample for multiple years and in both canopy and understory. Our results nevertheless indicate that species with longer forewings and with variable adult colourations had less variable abundance across the two study years.

## Supporting information

Data used in the study

## ACKNOWLEDGEMENTS

Fieldwork was carried out with the kind permission of the Forestry Experimental Station in Murowana Goślina (Poznań University of Life Sciences). Traps were generously lent by Martin Gossner and repaired and completed with advice from Edward Baraniak. We thank Jaroslaw Kordy, Öznur Tilkioğlu, Fatmanur Selvi, and Meriç Atar for help with setting moths, Pasquale Di Pratola for help with calculating phenology from the raw data, and Mariia Sokulska for measuring the moths. We thank Mike Shewring and anonymous reviewers for their insightful comments that helped improve the manuscript.

## Data Availability Statement

Data and code used in this manuscript will be made available from the Dryad Digital Repository and are here uploaded for review purposes.

## Author contributions

Freerk Molleman developed the initial concept of the study, participated in the fieldwork, set part of the moths, analysed the data, and wrote the first draft of the manuscript. Roman Wąsala identified the moths and compiled their diet, migration, and polymorphism information. Ahmet Tambay entered the data and contributed to the data analysis. Robert B. Davis performed comparative analyses, and Erki Õunap provided the phylogeny. Urszula Walczak participated in developing the concept, planning of the study, and in fieldwork. All authors contributed to the manuscript writing.

Larger moths showed less inter-annual abundance variation.

Those with adult colour variation also showed less inter-annual abundance variation. Species associated with oaks tended to have later abundance peaks.

